# Molecular Bases of the Membrane Association Mechanism Potentiating Antibiotic Resistance by New Delhi Metallo-*β*-Lactamase 1

**DOI:** 10.1101/2020.06.01.126664

**Authors:** Alessio Prunotto, Guillermo Bahr, Lisandro J. González, Alejandro J. Vila, Matteo Dal Peraro

## Abstract

Resistance to last-resort carbapenem antibiotics is an increasing threat to human health, as it critically limits therapeutic options. Metallo-*β*-lactamases are the largest family of carbapenemases, enzymes that inactivate these drugs. Among MBLs, New Delhi metallo-*β*-lactamase 1 has experienced the fastest and largest worldwide dissemination. This success has been attributed to the fact that NDM-1 is a lipidated protein anchored to the outer membrane of bacteria, while all other MBLs are soluble periplasmic enzymes. By means of a combined experimental and computational approach, we show that NDM-1 interacts with the surface of bacterial membranes in a stable, defined conformation, in which the active site is not occluded by the bilayer. Although the lipidation is required for a long-lasting interaction, the globular domain of NDM-1 is tuned to interact specifically with the outer bacterial membrane. In contrast, this affinity is not observed for VIM-2, a natively soluble MBL. Finally, we identify key residues involved in the membrane interaction of NDM-1, which constitute potential targets for developing therapeutic strategies able to combat resistance granted by this enzyme.

## INTRODUCTION

The ever-increasing prevalence of bacterial antibiotic resistance has led to a worldwide healthcare crisis ^1,2^. The emergence of multi-resistant and pan-resistant microbes, for which treatment options are very limited, is compounded by the lack of development of new antibiotics in the last decades ^3,4^. Amid calls from the WHO for urgent action in addressing this complex issue, estimates predict that infections from drug-resistant bacteria will become the major cause of death by 2050, if the current trend is not reversed^5^. The rise in carbapenem-resistant bacteria is particularly concerning, since this class of *β*-lactam antibiotics is reserved as a last resort option for life-threatening infections ^6,7^. The most prevalent cause of this resistance is the production of carbapenemases, hydrolases that degrade and inactivate these antibiotics ^8^. Metallo-*β*-lactamases (MBLs) are Zn(II)-dependent enzymes that represent the largest family of carbapenemases ^9,10^. MBLs can also hydrolyze other *β*-lactam antibiotics such as cephalosporins and penicillins ^11,12^. At the moment, there are no clinically approved inhibitors for MBLs, so that bacterial resistance mediated by these proteins cannot be countered ^13^.

New Delhi metallo-*β*-lactamase 1 (NDM-1) is a MBL first identified in 2008 ^14^ which has experienced a remarkably fast spread worldwide, having been detected in more than 100 countries distributed in all continents ^15,16^. This enzyme is not only widely disseminated in healthcare settings, but it also displays an unprecedented prevalence in the environment, with its coding gene being present in soil and water samples worldwide ^17,18^. The cellular localization of NDM-1 is unique among clinically relevant MBLs: while all others are soluble periplasmic enzymes, NDM-1 is a lipoprotein anchored to the inner leaflet of the outer membrane in Gram-negative bacteria ^19,20^. This post-translational modification is due to the presence of a lipidation signal within the signal peptide of the enzyme, which is recognized by a widely conserved lipoprotein biogenesis machinery located in the cell envelope of bacteria ^21,22^. The lipid moiety is covalently bound to a Cys residue (Cys26, **Figure 1a**) present in the signal peptide, which becomes the N-terminal residue of the mature protein, and is responsible for membrane anchoring.

**Figure 1.**
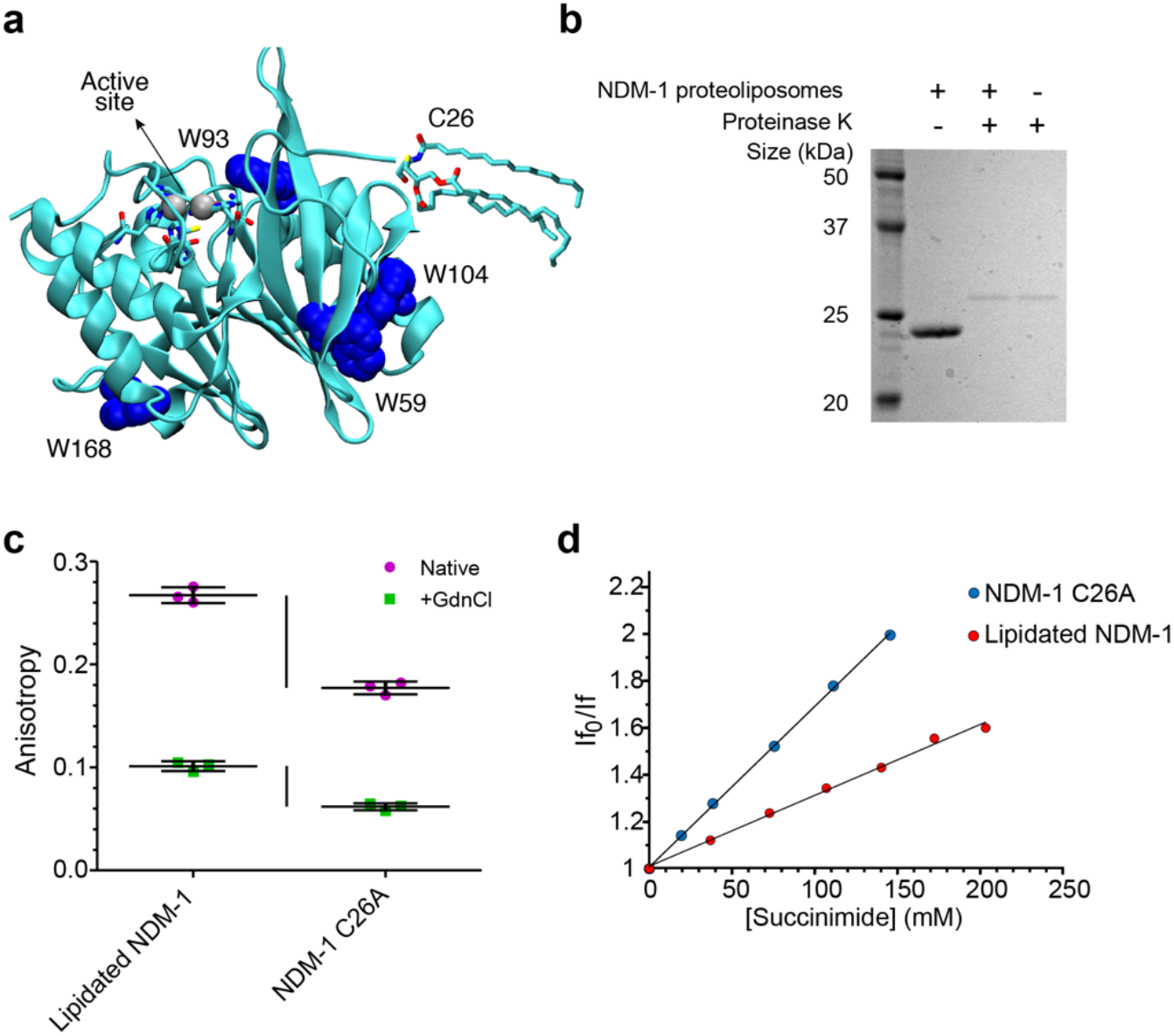
NDM-1 presents restricted mobility in contact with the membrane. (**a**) Location of tryptophan residues, Cys26 (highlighted in the lipidated form) and active site in the NDM-1 crystal structure (PDB: 5ZGE ^30^). (**b**) Analysis of NDM-1 orientation in proteoliposomes, using proteolysis by proteinase K. (**c**) Tryptophan fluorescence anisotropy values for lipidated NDM-1 in liposomes and soluble NDM-1 C26A, in absence (Native) or in presence (+GdnCl) of 4.5 M guanidinium chloride. Data for each protein are presented as mean ± st.dev. of 3 biological replicates, individual data points are presented in color. (**d**) Stern-Volmer plots of fluorescence quenching by succinimide of lipidated NDM-1 in liposomes and soluble NDM-1 C26A.

Membrane anchoring endows NDM-1 with unique features within the bacterial host that have favored its worldwide dissemination and persistence ^19^. First, this cellular localization stabilizes the enzyme upon conditions of metal starvation that occur at the infection sites. Second, and most important, membrane binding favors secretion of NDM-1 into outer-membrane vesicles (OMVs). These lipid-enclosed particles are released by all Gram-negative bacteria ^23^, and provide a mechanism for the dissemination of NDM-1. On one hand, these vesicles expand the spatial domain of antibiotic hydrolysis beyond the bacterial cells. On the other hand, OMVs also protect populations of bacteria that would otherwise be sensitive to antibiotics. Thus, membrane anchoring provides several evolutionary advantages to NDM-1.

The biochemical and biophysical characterization of NDM-1 has been restricted to its soluble domain. Indeed, all available crystal structures of the protein have been obtained with truncated soluble forms lacking the lipid group ^20,24,25^. In addition, computational studies on NDM-1 that are present in literature ^26–28^, generally focus on the catalytic action of the enzyme, regardless of its membrane anchoring. As a result, the molecular details of the interaction of NDM-1 with the membrane surface are unknown. There are several unaddressed questions regarding the membrane association mechanism of NDM-1: (1) how does the soluble domain of NDM-1 (in particular, the active site) orient with respect to the bacterial membrane? (2) is lipidation the only mechanism contributing to membrane association, or are there specific interactions of the bilayer with the soluble domain? (3) is NDM-1 tuned to interact more favorably with the outer membrane environment rather than with the inner membrane? Answering these questions could assist in designing drugs that thwart these interactions, thus hampering the distinct advantages conferred by membrane anchoring.

Here, by combining experimental and computational approaches, we show that NDM-1 adopts a defined orientation with respect to the membrane, characterized by the synergistic anchoring action of the lipidated cysteine and a molecular surface with specific affinity for the outer bacterial membrane. As a result, the NDM-1 active site faces the periplasmic space and is exposed to the solvent, hence without spatial restrictions that limit its hydrolytic action. We have identified residues that are critical for this interaction, as well as the effect of the membrane composition on the protein-membrane interaction. Overall, this picture reveals that the soluble domain contributes favorably to the interaction with the outer membrane of Gram negative bacteria, potentiating the effect of enzyme lipidation.

## RESULTS AND DISCUSSION

### NDM-1 presents restricted mobility in contact with the membrane

As a first step to characterize the interaction of NDM-1 with the bacterial membrane, we aimed to obtain information about the proximity and relative mobility of NDM-1 in its anchored form with respect to the membrane bilayer surface. We decided to use proteoliposomes as a model system amenable to in vitro characterization. Purified NDM-1 was introduced into liposomes with a phospholipid composition mimicking the inner leaflet of the outer membrane of *E. coli*, which is formed by neutral Phosphatidyl-Ethanolamines (PEs, 91%), and by two anionic lipids: Cardiolipins (CDLs, 6%) and PhosphatidylGlycerols (PGs, 3%) ^29^. This was achieved by using liposomes with 91% POPE (1-palmitoyl-2-oleoyl-phosphatidylethanolamine), 6% tetraoleoyl cardiolipin, and 3% POPG (1-palmitoyl-2-oleoyl-phosphatidylglycerol). NDM-1 bound to proteoliposomes may be oriented outwards (with the lipid group inserted within the outer leaflet of the membrane), inwards (with the protein encapsulated within the liposome), or with the presence of mixed populations exhibiting each orientation. Treatment of proteoliposomes with proteinase K resulted in full degradation of NDM-1 (**Figure 1b**), indicating that all protein molecules were located on the outer surface of the liposomes.

We then sought to evaluate whether NDM-1 assumes a fixed orientation with respect to the membrane or if it is sampling different conformations. Fluorescence anisotropy measurements enable determining the tumbling rates of molecules in solution, as higher rotational correlation times cause increased anisotropy in the emitted fluorescence. We measured the fluorescence anisotropies of lipidated NDM-1 in liposomes, and of the soluble form of the enzyme, NDM-1 C26A. In this mutant, the lipidated Cys26 residue is replaced by an alanine, resulting in a soluble periplasmic enzyme ^19^. Since NDM-1 possesses four tryptophan residues distributed along the protein structure (**Figure 1a**), we exploited the intrinsic anisotropy of the protein upon excitation at 298 nm. The membrane-anchored form of the protein presents a much higher (0.267 ± 0.008) anisotropy than the soluble NDM-1 C26A variant (0.177 ± 0.006), suggesting that NDM-1 assumes a fixed orientation with respect to the membrane (**Figure 1c**). A different scenario was found when measuring fluorescence anisotropy of the unfolded states of both variants (by addition of 4.5 M guanidinium chloride). Indeed, the difference in anisotropy between lipidated and soluble NDM-1 was much smaller compared to the folded proteins (0.039 vs. 0.090) (**Figure 1c**). Thus, despite the proximity to the liposome indeed induces a restricted mobility, the soluble domain seems to have a specific interaction with the bilayer that is only present in the folded protein.

Since three out of the four Trp residues in NDM-1 are partially exposed to the solvent (**Figure 1a**), we sought to confirm that the protein contacts the membrane surface by determining if any of these residues becomes occluded in the anchored protein. We carried out Trp fluorescence quenching assays with lipidated NDM-1 in liposomes and soluble NDM-1 C26A, using the non-ionic quencher succinimide. Since fluorescence quenching requires almost direct contact of the fluorophore and quencher, if any of the Trp residues is directly contacting the surface of the membrane, it should change its accessibility to the quencher in the membrane bound form of NDM-1 with respect to NDM-1 C26A in solution. Quenching data fit to a linear Stern-Volmer plot, indicating that all Trp residues both in NDM-1 C26A and in the liposome-bound NDM-1 are susceptible to quenching by succinimide. However, the two samples presented differences, as the lipidated protein displayed a significantly smaller Stern-Volmer constant (KSV) with respect to the soluble form (**Figure 1d**). A smaller KSV correlates to a restricted diffusion constant of the quencher towards the protein. As a consequence, the same degree of quenching in the lipidated protein can be achieved with a higher concentration than that required for the soluble NDM-1 C26A. We attribute this finding to the proximity of the liposome in the case of lipidated NDM-1.

In summary, fluorescence anisotropy and fluorescence quenching experiments point to a close interaction of NDM-1 with the membrane surface, in which the protein adopts a stable conformation with respect to the bilayer.

### The soluble domain of NDM-1 contributes to membrane association

In order to characterize in further detail the interaction of NDM-1 with the membrane, we conducted coarse-grained (CG) molecular dynamics (MD) simulations using the Martini ^31^ force field. We generated models of NDM-1 in the lipidated (wild type) and soluble forms (C26A), considering both their holo and apo forms, i.e. with and without zinc ions at the catalytic site (**Figure S1a**). The membrane bilayer was modeled with the same lipid composition used in the liposomes mimicking the inner leaflet of the bacterial outer membrane.

The enzyme was located at 4 nm at least from the bilayer in the starting geometry in order to avoid initial protein-membrane interactions that may bias the simulations. After exploring several conformations, lipidated NDM-1 spontaneously binds to the membrane by insertion of the lipid moiety into the bilayer. NDM-1 C26A also reaches a stable binding conformation to the membrane within the same timescale (**Figure 2a,b**), showing a similar number of binding events compared to the lipidated variant (**Figure 2c**). Indeed, we observed membrane binding in all the CG-MD replicas for both protein forms. The membrane-bound forms, which adopt similar orientations in all the different replicas, reveal that the active site is not occluded by the interaction with the membrane (see following section). Remarkably, the soluble and lipidated NDM-1 display the same surface of interaction with the membrane. The CG-MD trajectories show that the tip of the N-terminal tail has a natural propensity to interact with the membrane even in absence of lipidation.

**Figure 2.**
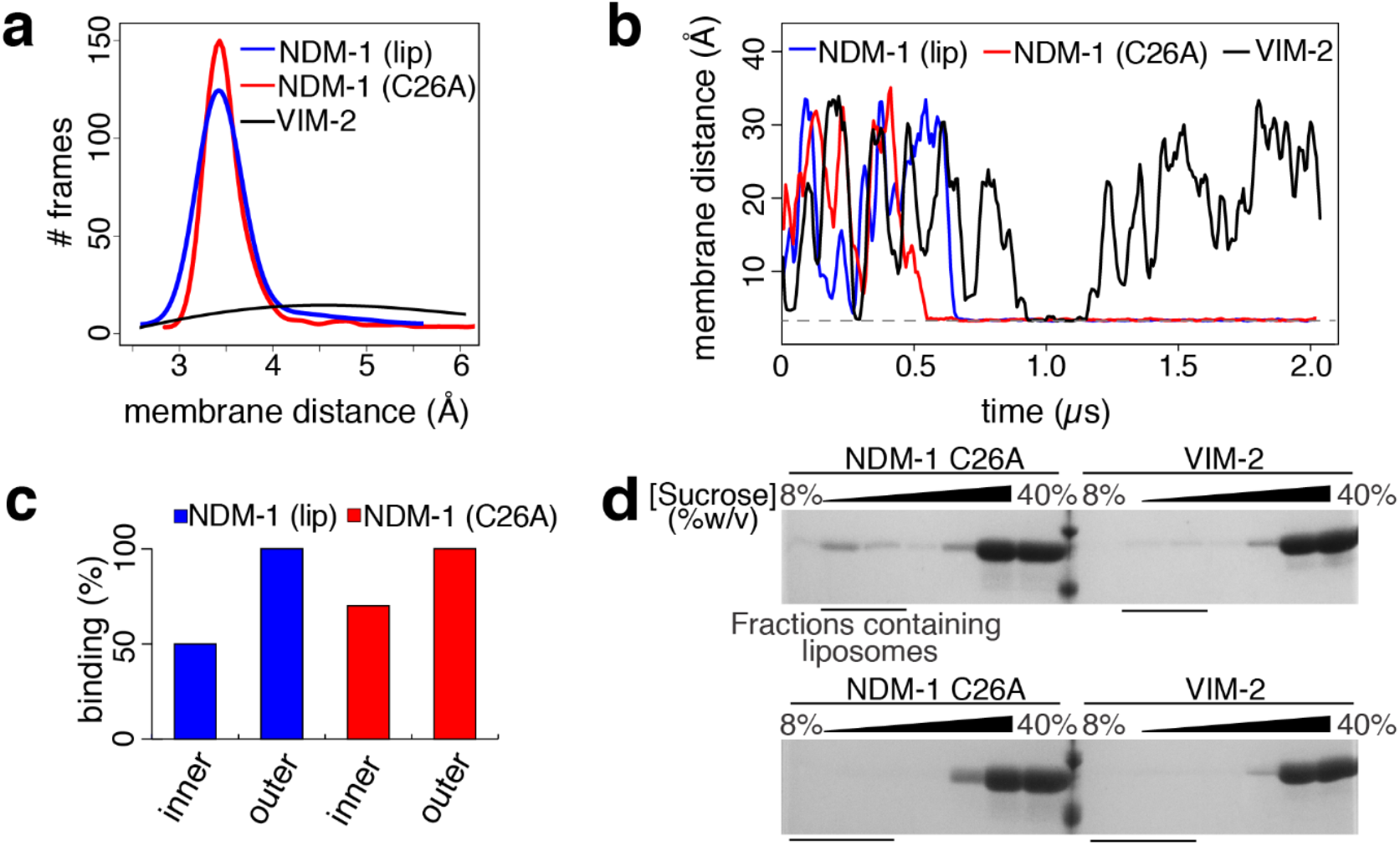
The soluble domain of NDM-1 contributes to membrane association. (**a**) Distribution of CG-MD trajectory frames, collected every 750 ps, for to the protein/membrane distance of lipidated NDM-1, soluble NDM-1 (NDM-1 C26A) and VIM-2. (**b**) Protein-membrane distance evolution for lipidated and soluble NDM-1, and for VIM-2. One representative replica for each protein is reported. (**c**) Percentage of binding events observed during the CG MD simulations for the tested systems. (**d**) SDS-PAGE analysis of sucrose gradient fractions from liposome flotation assays of NDM-1 C26A and VIM-2. The flotation assays were carried out using liposomes made with an outer membrane composition (top) or *E. coli* Polar Lipid Extract (bottom).

The Zn(II) content at the active site does not affect the interaction with the membrane, as we observed a similar binding propensity in both the apo and the holo form, (**Table S1a**) despite their different net charges. This can be attributed to the fact that the active site is oriented on the opposite side of the surface anchored to the membrane. Additionally, the Zn(II) ions are not located at the protein surface, reducing their impact in surface electrostatics.

These results suggest that NDM-1 has the tendency to interact with the outer bacterial membrane regardless of its metal content and lipidation at Cys26. Despite lipidation is fundamental for membrane association and *in vivo* requires a complex biogenesis machinery ^21^, these simulations suggest that the soluble domain of NDM-1 has native electrostatic features that favor its interaction with the membrane bilayer. The effect of lipidation probably acts on reinforcing and stabilizing membrane anchoring on a longer time scale.

We also simulated the enzyme VIM-2 as a model of a soluble periplasmic MBL in its native state. In this case, no interaction with the membrane was observed (not even transient, **Figure 2a,b** and **Table S1b**), in contrast to soluble NDM-1 C26A. It appears that different MBLs with high degree of structural homology can present distinct affinities towards the bacterial membrane. Therefore, we can conclude that the soluble domain of NDM-1 is specifically adapted to interact with the membrane, in contrast to VIM-2.

To test the predictions of the molecular simulations, we carried out liposome flotation assays by incubating NDM-1 C26A with liposomes mimicking the outer membrane composition of *E. coli*. Samples were loaded at the bottom of a discontinuous sucrose gradient and ultracentrifuged. Under these conditions the liposomes tend to float along the gradient towards lower sucrose concentrations, carrying with them bound proteins, while free protein remains at the bottom of the gradient. We used SDS-PAGE to analyze the distribution of NDM-1 C26A after ultracentrifugation, and clearly observed that this soluble enzyme indeed binds the vesicles (**Figure 2d**, top), with a portion of the protein having migrated with the liposomes to a lower sucrose concentration. Instead, no binding of VIM-2 could be detected (**Figure 2d**, top).

These results confirm that NDM-1 has the ability to bind to the membrane regardless of the presence of the lipidated cysteine residue. The low amount of bound NDM-1 that we observed suggests a transient interaction, leading to a high proportion of the protein being located in its initial position at the bottom of the gradient. The presence of the lipid group in the wild-type protein is required to fully stabilize the interaction, granting a more defined and long-lasting contact between the protein and the membrane.

In order to explore the effect of membrane localization and composition, we used CG-MD simulations to probe NDM-1 association to the inner bacterial membrane. In *E. coli*, the inner bacterial membrane is characterized by a significantly higher content in charged lipids, in particular PGs (67% PEs, 28% PGs, and 5% CDLs) ^32^. The simulations revealed that NDM-1 has a lower affinity for the inner membrane compared to the outer bilayer: the binding events of all protein states (i.e., apo/holo form and with/without lipidation) represent 60% of the CG-MD replicas, in contrast to the 100% of replicas showing binding events to the outer membrane (**Figure 2c**, **Table S1a**, **Figure S1b**). Analogous simulations on the soluble MBL VIM-2 revealed no affinity for the inner bacterial membrane model.

These predictions were also validated by performing liposome flotation assays for NDM-1 C26A with liposomes produced with the *E. coli* polar lipid extract, which contains a lipid composition mimicking the inner membrane, resulting in no detectable binding (**Figure 2d**, bottom). Similar experiments with VIM-2 also resulted in no binding, confirming the behavior predicted by CG-MD simulations. Based on these results, we then decided to identify the molecular features favoring the interaction of NDM-1 with the outer bacterial membrane.

### Specific interaction of the soluble domain of NDM-1 with the outer membrane

The analysis of the CG-MD trajectories reveals that NDM-1 interacts with the membrane by means of a specific patch at the protein surface. During the initial stages of the simulation, NDM-1 explores different orientations, but only one of them elicits a neat binding event (**Figure 3a**). While the lipidated Cys26 inserts directly into the membrane, the globular domain interacts with the membrane by means of the β-strand domain spanning from Thr41 to Val58 and the segment between Asn103 and Pro112 (which includes a portion of α-helix, Asn103-Ile109, and a portion of a loop, Asn110-Pro112, **Figure 3a**). These two regions include six charged residues: Arg45, Arg52 and Lys106, Asp43, Asp48 and Glu108 (**Figure S1c).** Particularly the three basic amino acids might be key residues in providing the affinity of the globular domain of NDM-1 towards the membrane. While the overall charge of NDM-1 is negative (−5e in the holo form and −7e in the apo form), these positively charged residues could provide a favorable electrostatic interaction with the membrane. It is important to observe that the identified configuration orients the active site of NDM-1 exposed to the solvent: the enzyme is therefore still able to capture, host and cleave β-lactam molecules, without being hindered by the presence of the membrane. In addition, we analyzed the variation in the tilt angle of the protein across the simulations, which is the angle formed by the catalytic site, the center of mass of the protein, and its projection on the membrane. NDM-1 adopted an orientation with a constant tilt angle (~150°, **Figure 3b**), reflecting that the protein adopts a well-defined conformation with respect to the membrane surface.

**Figure 3.**
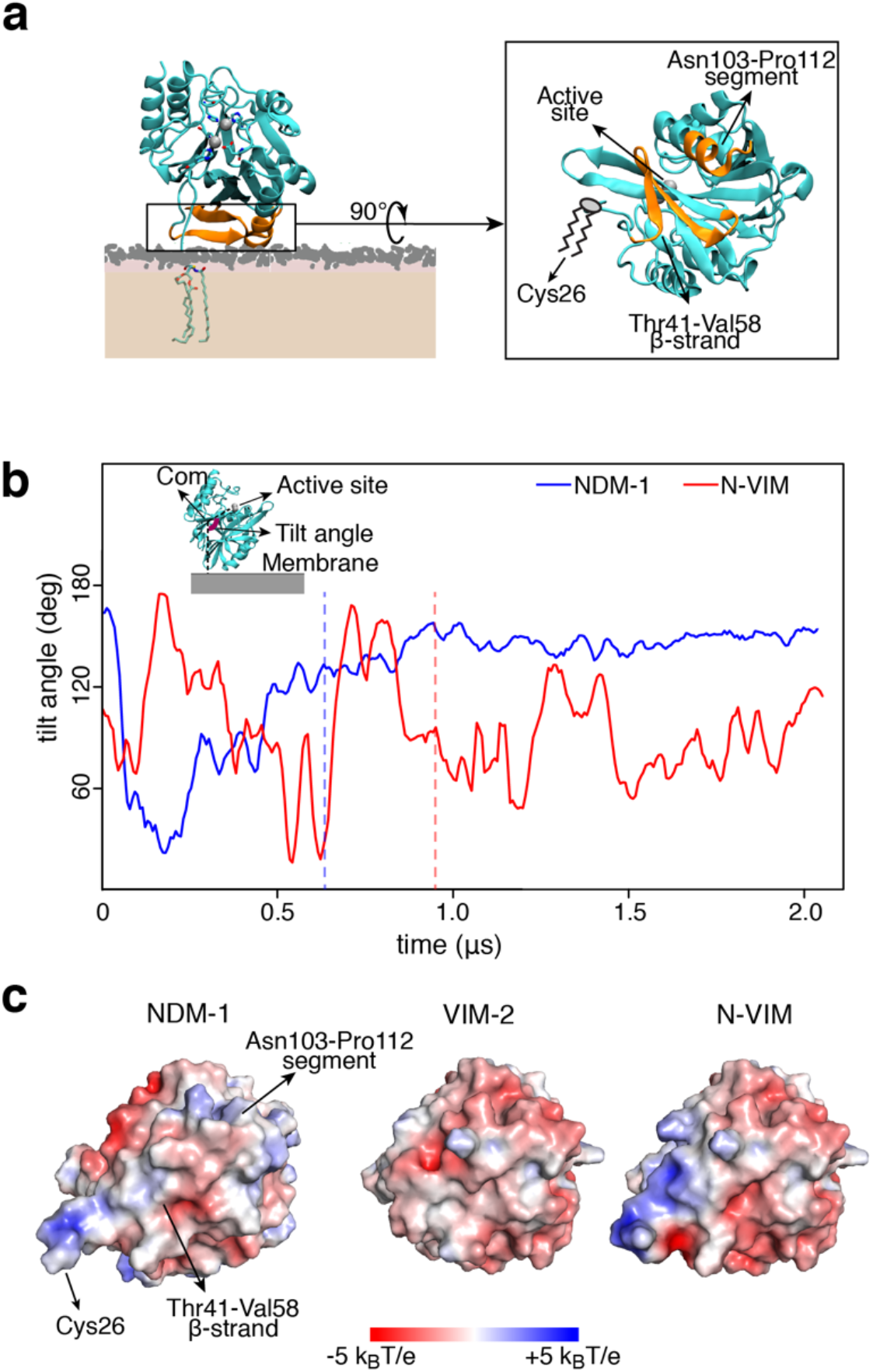
The soluble domain of NDM-1 is optimized to interact with the outer bacterial membrane. (**a**) Final and stable orientation of NDM-1 with respect to the membrane, and protein domains that are involved. The active site is exposed to the solvent and not hindered by the membrane. (**b**) Tilt angle definition and evolution of the tilt angle vs. time, for NDM-1 and N-VIM at the outer bacterial membrane. The tilt angle is defined as the angle formed by the active site, the center of mass of the protein and the projection of the center of mass of the protein on the plane of the membrane. The vertical dashed lines represent the moment in which the anchoring occurs. (**c**) Electrostatic potential of the protein-membrane surface of interaction (seen from the membrane surface) for NDM-1, VIM-2 and N-VIM.

To further evaluate the contribution of the soluble domain to this interaction, we tested the behavior of a chimera between NDM-1 and VIM-2. This protein, named N-VIM, contains the main core of the soluble enzyme VIM-2 with the N-terminal region of NDM-1 (including the lipidation site Cys26), resulting in a membrane-bound protein ^19^. In the CG-MD simulations, the chimeric protein N-VIM did not show a significantly improved binding with respect to the native, soluble VIM-2 protein (**Table S1c**). Analysis of the tilt angle shows that N-VIM did not achieve a stable membrane anchoring, in contrast with wild type NDM-1 (**Figure 3b**). Clustering analysis of the MD trajectories confirms that the orientation of N-VIM with respect to the membrane covers a broad range of conformations, in contrast to the stable and defined interaction observed for NDM-1 (**Figure S1d**).

The diverse interaction of NDM-1, VIM-2 and N-VIM can be accounted for by analyzing the electrostatic potential of the 3 enzymes (**Figure 3c**). NDM-1 shows a larger surface of positive electrostatic potential with respect to VIM-2 in the region that drives membrane anchoring to the negatively charged phospholipid bilayer. Comparing the interaction surface of these proteins, we observed that VIM-2 is almost completely negatively charged, whilst N-VIM is characterized by an electrostatic profile that is intermediate between NDM-1 and VIM-2 (**Figure S1e**). The behavior of N-VIM is probably caused by the lack of most of the surface of interaction identified in NDM-1. In particular, the surface of NDM-1 is more positive than N-VIM in the region corresponding to the Asn103-Pro112 segment, which is reasonable since in this area N-VIM has the same primary sequence as VIM-2.

In order to better define the molecular interaction at the protein-membrane interface, we performed all-atom (AA) MD simulations based on the CG models. These calculations confirmed the affinity between NDM-1 and the bacterial membrane. The behavior of NDM-1 largely coincided in both types of simulations, although we observed a partial reorientation of the protein on the membrane surface (**Figure 4a**). This suggests a more prominent role for the β-strand Thr41-Val58, with respect to the segment Asn103-Pro112, as confirmed by the analysis of the residues contacts (**Figure S2a**). Despite these minor differences, both CG- and AA-MD simulations reveal that NDM-1 adopts a stable orientation with respect to the membrane, in excellent agreement with the fluorescence anisotropy results (**Figure 1**).

**Figure 4.**
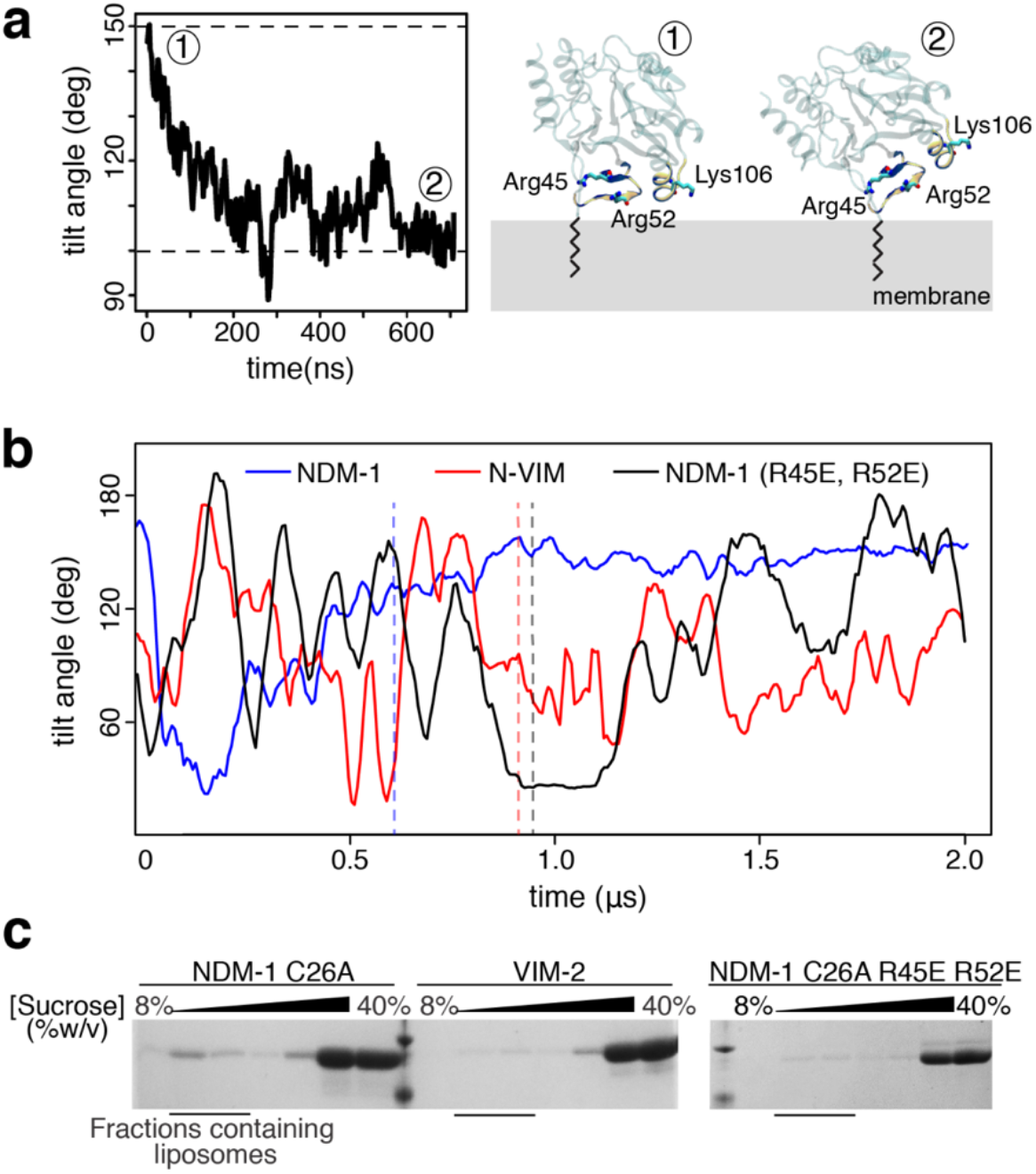
Two positively charged residues drive the interaction between NDM-1 and the bacterial membrane. (**a**) Tilt angle evolution of NDM-1 during all-atom MD simulations (left) and comparison between the surface of interaction identified during the CG (~150°) and all-atom (~100°) MD simulations (right). (**b**) Comparison of tilt angle evolution during MD simulations for wild type NDM-1, N-VIM and mutant NDM-1 R45E-R52E. The vertical dashed lines represent the moment in which the membrane anchoring occurs. (**c**) SDS-PAGE analysis of sucrose gradient fractions from liposome flotation assays of NDM-1 C26A, VIM-2 and NDM-1 C26A R45E-R52E, using liposomes with an outer membrane composition.

These simulations also agree in pinpointing the Thr41-Val58 stretch as the main contributor to the interaction between NDM-1 and the outer membrane. To challenge this hypothesis, we ran CG-MD simulations of an NDM-1 variant in which the positively charged residues Arg45 and Arg52 were replaced by glutamates. This *in silico* mutation resulted in a reduction of the total number of binding events to the outer membrane with respect to wild type NDM-1, that dropped from 100% to 20% (**Table S1a**). The tilt angle of the bound NDM-1 double mutant R45E-R52E resembled that of N-VIM (**Figure 4b**), suggesting that these mutations indeed destabilize the protein-membrane interaction.

To validate these predictions, we experimentally evaluated the impact of these mutations in the interaction of the soluble form of the protein with the membrane. Flotation assays with liposomes mimicking the outer membrane showed that these mutations indeed reduced the binding of NDM-1 to the liposomes, with a behavior closer to that of VIM-2 than to wild type NDM-1 (**Figure 4c**). Overall, these data provide compelling evidence of the role of the electrostatic interaction between the soluble domain of NDM-1 and the outer membrane in maintaining membrane association. We conclude that the soluble domain of NDM-1, in contrast to VIM-2, has been selected during evolution to better interact with the membrane, and that the presence of the N-terminal lipid group is not sufficient on its own for a stable interaction with the membrane.

### Role of cardiolipin in NDM-1 binding to the outer bacterial membrane

CG-MD simulations showed a clear preference of NDM-1 for the outer bacterial membrane model compared to the inner one (100% vs. 60% of replicas showing binding, **Figure 2c**, **Figure S1b** and **Table S1a**), despite its lower content in charged lipids (9% vs. 33% of anionic lipids). This is against the contention that electrostatics is expected to be the main driving force of the interaction between phospholipid bilayers and peripheral binding proteins. The role of charged lipids, and cardiolipins (CDLs) in particular, has been reported as fundamental for the correct functioning of bacterial and mitochondrial membrane proteins ^33–36^. In order to evaluate the role of CDLs in the interaction between the bacterial membrane and NDM-1, we conducted CG-MD simulations in absence of this specific lipid, but keeping fixed the amount of PGs. This resulted in a significant reduction in the number of binding events (40% of replicas compared to 100% in presence of CDLs, **Table S1a**). The same trend was observed in simulations performed with an inner bacterial membrane model without CDLs (**Table S1a**). In both cases, NDM-1 approaches and explores the bilayer surface without ever establishing a stable contact (**Figure 5a**), resembling the situation previously described for VIM-2. These findings support the idea that the nature of the lipids present in the membrane plays also a fundamental role in NDM-1 recognition and association.

**Figure 5.**
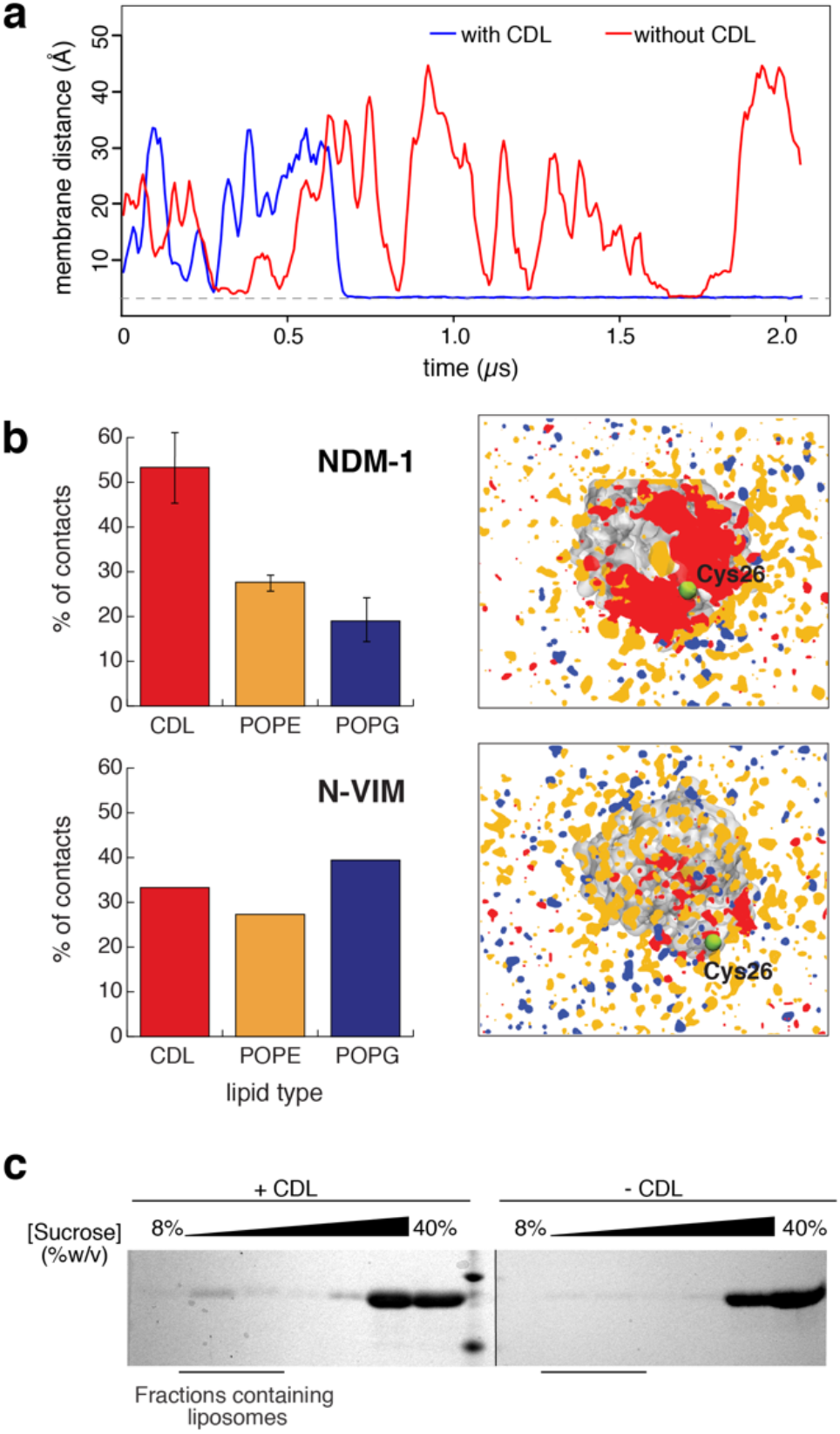
Role of cardiolipin in NDM-1 binding to the outer bacterial membrane. (**a**) Distance evolution of NDM-1 vs. outer membrane distance during CG MD simulations, with and without CDLs. One representative replica for each membrane model is reported. (**b**) Lipid contacts between the enzymes and the different lipid types that compose the bilayer, normalized according to lipid composition of the membrane (left); lipid distribution in proximity of NDM-1 and N-VIM, averaged over all the MD trajectories (right); (**c**) SDS-PAGE analysis of sucrose gradient fractions from liposome flotation assays of NDM-1 C26A, using liposomes with an outer membrane composition (+CDL) or liposomes with an equivalent composition but lacking cardiolipin (−CDL).

Driven by these results, we further analyzed the average occupancy of lipids in proximity of membrane-bound NDM-1. Clusters of charged lipids (in particular CDLs) form close to the NDM-1 binding surface (**Figure 5b**). In contrast, it was not possible to identify a specific pattern of lipid contacts for VIM-2, since in this case the contacts are transient and never turn into stable and long-term interactions. However, the lipid arrangement within the membrane patch seems to be completely non-specific (**Figure S2b**). This suggests that the presence of NDM-1 in close proximity with the membrane is fundamental in order to drive the assembly of lipid clusters.

In order to further understand whether the physical proximity to the membrane (regardless of a specific affinity) could drive the recruitment of charged lipids, we tested the behavior of N-VIM. In the only replica in which membrane binding occurred for N-VIM, we could not detect any specific, more dominant, contact with lipid species (**Figure 5b**). We conclude that recruitment of charged lipids – specifically CDLs – is a fundamental step for the generation of a stable and strong association with the membrane, which needs to be driven by a specific protein surface. Lipid distribution analysis on the AA-MD trajectories indicates that clusters of CDLs are still present and contribute to maintain a strong contact between NDM-1 and the phospholipid bilayer (**Figure S2c**). In particular, we observed two CDL molecules generating long-lasting interactions (i.e. for the whole length of the simulation) with the residues that we identified in the previous section, i.e. Arg45 and Arg52 (**Figure S2d**). The proximity to CDLs confirms the importance of these aminoacids in generating a strong and stable contact with the membrane, and explains why the NDM-1 R45E-R52E mutant is characterized by a smaller interaction with the bilayer (**Figure 4b,c**).

To experimentally test these predictions, we carried out liposome flotation assays with NDM-1 C26A using liposomes containing 97% PE and 3% PG (without CDLs). Removal of CDLs indeed abolishes binding of soluble NDM-1 to the liposomes (**Figure 5c**), in agreement with the weaker interaction predicted by simulation studies.

These results pinpoint a crucial role of CDLs in the recruitment and binding of NDM-1 to the membrane, confirming both the impact of the membrane composition for membrane-protein association and the role of electrostatics in the interaction of the soluble domain of NDM-1 to the membrane.

## CONCLUSIONS

NDM-1 has been one of the major molecular causes of carbapenem resistance in several outbreaks of opportunistic and pathogenic bacteria ^37^. The hydrolytic abilities of this enzyme are similar to those of other clinically relevant MBLs ^14^, and cannot explain its success as a resistance determinant. The unique worldwide dissemination of the zinc-dependent lactamase NDM-1 has been attributed to the fact that this enzyme is bound to the inner leaflet of the outer membrane of Gram-negative bacteria ^19^, in contrast with the rest of MBLs that are soluble, periplasmic proteins.

Lipidation allows membrane anchoring of NDM-1, being a unique characteristic of this enzyme. In the present work, we have characterized the mechanistic details of the association of NDM-1 with the bacterial membrane, demonstrating that the enzyme maintains a stable orientation with respect to the bilayer, while exposing its active site to the solvent, therefore preserving its catalytic activity. A defined protein surface contacts the membrane and contributes to the strong electrostatic interaction that acts synergistically with lipidation at Cys26. These results are strongly supported by molecular simulations at different levels of resolution, as well as by fluorescence studies and liposome flotation assays in a model mimicking the outer bacterial membrane. This membrane-protein interaction is characteristic of NDM-1, since in the case of the soluble MBL VIM-2, no interactions with the membrane are predicted by the simulations nor observed in the interaction with liposomes. Instead, a soluble version of NDM-1 is still able to interact with the membrane despite lacking the lipid moiety,

These findings allow us to conclude that, while the lipid group attached to the N-terminal Cys26 of NDM-1 is of crucial importance in maintaining membrane association, the globular domain of the protein possesses a native affinity towards the bacterial membrane. This phenomenon has been reported for other peripheral membrane proteins ^35,38^. The presence of the lipid group is essential in order to stabilize the protein/membrane interaction on a long-term timescale, resulting thus in robust anchoring. Lipidation by itself is not sufficient, since the chimeric protein N-VIM (a membrane-anchored version of VIM-2) has a poorer membrane association than NDM-1. Both the lipidation and the recruitment of CDLs are essential for generating a strong and stable anchoring to the membrane. The anchoring mechanism of NDM-1 is therefore specific for the outer bacterial membrane, and is strongly compromised in presence of inner bacterial membrane or CDL-free bilayers. Thus, evolution has optimized the ability of NDM-1 to interact with the outer membrane milieu that is determined by its lipidation signal and the bacterial lipoprotein biogenesis machinery.

We have identified a positively charged patch in the surface of NDM-1 that is responsible for this additional mechanism of association. However, the presence of positively charged residues does not seem to be the only responsible for driving strong and stable association between the protein and the membrane. Indeed, the positive residues must pack in a proper way, in order to be able to attract specific clusters of lipids (such as CDLs) that stabilize the interaction with the protein. The induction of lipid clusters formation by the protein is crucial to guarantee a long-lasting interaction with the bilayer, as suggested by the CG-MD simulations of the N-VIM chimera, which is not able to attract CDL aggregates.

Finally, this contact surface contains two basic residues (i.e., Arg45 and Arg52) that are important in defining a stable interaction with the bilayer, which is disrupted by the R45E-R52E substitutions. Notably, these two Arg residues are conserved in all reported NDM variants (28 so far). The first 16 NDM variants have been shown to be bound to the membrane ^39^. While the other 12 still lack a biochemical characterization, all variants have a conserved lipobox in the signal peptide, suggesting that all of them are effectively membrane proteins. Thus, the conservation of the two positively charged Arg residues confirms the essentiality of this patch for the membrane-protein interaction, and reveal that the globular domain of NDM variants has been selected by evolution with features that favor its cellular localization, and ultimately, secretion to vesicles. New therapeutic strategies targeting these positions could inhibit the association of NDM variants with the membrane surface, aiding in the fight against this resistance determinant by negating the advantages conferred by NDM unique cellular localization.

## MATERIALS AND METHODS

### Cloning and generation of MBL mutants

The full length NDM-1 gene was amplified by PCR from the pMBLe NDM-1 plasmid ^19^ using the NDM-1 NdeI Fw (5’ TATACATATGGAATTGCCCAATATTATGCACC 3’) and NDM-1-TEV-TwinST-HindIII Rv1 (5’ GAACCACCACCCTTTTCGAATTGTGGGTGAGACCAGCCCTGAAAATACAGGTTTTCGCGCAGCTTGTCGGCCATGC 3’) primers. This product was then used as a template for another round of PCR, using the NDM-1 NdeI Fw and NDM1-TwinST-HindIII Rv2 (5’ GACGTAAGCTTCTACTTTTCGAATTGTGGGTGAGACCACGCAGAACCACCAGAACCACCACCAGAACCACCACCCTT TTCG 3’) primers. The product was cloned into the NdeI and HindIII sites of the pET-26 vector, obtaining the pET-26 NDM-1 TEV TST plasmid. From this plasmid, the pET-26 NDM-1 C26A TEV TST plasmid was obtained through site directed mutagenesis by plasmid amplification as previously described ^40^, using primers C26A Fw (5’ CATTGATGCTGAGCGGGGCGATGCCCGGTGAAATC 3’) and C26A Rv (5’ GATTTCACCGGGCATCGCCCCGCTCAGCATCAATG 3’).

### Liposome preparation

Pure lyophilized phospholipids (1-palmitoyl-2-oleoyl-phosphatidylethanolamine, tetraoleoyl cardiolipin, and 1-palmitoyl-2-oleoyl-phosphatidylglycerol) and *E. coli* Polar Lipid Extract were purchased from Avanti Polar Lipids. The lipids were dissolved in chloroform and after mixing the required proportions of each pure lipid, the lipid mixtures were dried under a nitrogen atmosphere and then kept under vacuum for 2h. The dried lipid film was hydrated with 50 mM HEPES pH 7, and heated at 65°C for 1h with periodic vortexing. Lipid suspensions were frozen in liquid nitrogen and then thawed at 65°C, for a total of 5 cycles, and afterwards were passed through a 400 nm polycarbonate filter using an Avanti Miniextruder apparatus (Avanti Polar Lipids) at 65°C, with >20 passes through the device.

### Purification of lipidated NDM-1 and soluble NDM-1 C26A

Full length NDM-1, including its signal peptide and lipidation signal, was overexpressed in *E. coli* BL21 (DE3) cells using the pET-26 NDM-1 TEV TST plasmid. The NDM-1 protein is produced from this vector with a fusion to its C-terminus of a TEV protease cleavage site followed by the Twin-StrepTag peptide ^41^, which allows affinity purification by binding to a StrepTactin Sepharose resin (GE Healthcare). Cells were grown in LB medium at 37°C with agitation to OD600nm = 0.8, and protein expression was induced by addition of 0.5 mM IPTG. Cultures were then grown at 20°C for 16 h. The cells were collected by centrifugation, resuspended in 50 mM HEPES pH 7.5, 200 mM NaCl, and ruptured at 15000 psi using an Avestin Emulsiflex C3 high pressure homogenizer. Cell debris was spun down by centrifugation at 14000g and 4°C for 20 min, and membranes were isolated by ultracentrifugation for 1h at 4°C and 125000 g in a Beckman SW Ti90 rotor. The lipid-anchored protein produced in this way was extracted from membranes by solubilization with 1% w/v of the nonionic detergent Triton X-100 in 50 mM HEPES pH 7.5 200 mM NaCl, followed by ultracentrifugation using the same conditions as before to remove non-solubilized material. Protein was purified by affinity chromatography with StrepTactin Sepharose resin, eluted with 2.5 mM desthiobiotin in 10 mM HEPES pH 7.5 200 mM NaCl 0.033 % w/v Triton X-100, and the affinity tag was removed by cleavage with TEV protease. The protein was dialyzed for 16h versus >100 volumes of 10 mM HEPES pH 7.5 200 mM NaCl 0.033 % w/v Triton X-100, 0.6 mM βME, and then for 4h versus >100 volumes of 10 mM HEPES pH 7.5 200 mM NaCl 0.033 % w/v Triton X-100, ZnSO4 0.1 mM. Finally, the protein was passed a second time through the StrepTactin Sepharose resin to remove the residual non-processed protein.

Proteoliposomes containing NDM-1 were then prepared by incubation of the purified protein with preformed liposomes and 0.7% w/v Triton X-100, and detergent removal was carried out with SM2 Bio-beads (Biorad). Proteoliposomes were separated from non-incorporated protein by ultracentrifugation on a discontinuous sucrose gradient (8% w/v, 25% w/v and 55% w/v), with protein incorporation into proteoliposomes being around 25% of the total NDM-1 added.

NDM-1 C26A was purified from *E. coli* OverExpress C43 (DE3) cells transformed with the pET-26 NDM-1 C26A TEV TST plasmid. Cells were grown in LB medium at 37°C with agitation to OD600nm = 0.8, and protein expression was induced by addition of 0.5 mM IPTG. Cultures were then grown at 20°C for 16 h. The cells were collected by centrifugation and resuspended in 50 mM HEPES pH 7.5, 200 mM NaCl. The cells were then ruptured at 15000 psi using an Avestin Emulsiflex C3 high pressure homogenizer, and cell debris was spun down by centrifugation at 14000g and 4°C for 20 min. Protein was purified by affinity chromatography with StrepTactin Sepharose resin, eluted with 2.5 mM desthiobiotin in 10 mM HEPES pH 7.5 200 mM NaCl, and the affinity tag was removed by cleavage with TEV protease. The protein was dialyzed for 16h versus >100 volumes of 10 mM HEPES pH 7.5 200 mM NaCl 0.033 % w/v Triton X-100, 0.6 mM βME, and then for 4h versus >100 volumes of 10 mM HEPES pH 7.5 200 mM NaCl 0.033 % w/v Triton X-100, ZnSO4 0.1 mM. Finally, the protein was passed a second time through the StrepTactin Sepharose resin to remove the residual non-processed protein.

### Fluorescence anisotropy determinations

Determinations of tryptophan fluorescence anisotropy were carried out in a Varian Cary Eclipse Spectrofluorometer using a 10×2 mm optical path quartz cuvette. Sample excitation was performed at 298 nm, to take advantage of the higher fundamental anisotropy of tryptophan at this wavelength, and fluorescence emission was collected at 370 nm. The excitation and emission slit widths were set to 5 nm and 10 nm, respectively, and a 360-1100 nm band pass filter was used in the emission optical path to remove scattered light. An integration time of 5 s was used for each measured fluorescence signal, and 5 replicate anisotropy determinations were acquired for each sample and averaged. To evaluate whether there is an appreciable contribution from light scattering to the anisotropy values previously determined, we added an equivalent amount of liposomes (not containing any protein) to samples of NDM-1 C26A. The observed anisotropies were similar to those of the soluble protein in absence of liposomes, indicating no interference from scattered light in our analysis.

Fluorescence quenching. Tryptophan fluorescence quenching determinations were carried out in a Varian Cary Eclipse Spectrofluorometer using a 5×5 mm optical path quartz cuvette. Sample excitation was performed at 280 nm, and fluorescence emission was collected from 300 to 450 nm, at 60 nm/s scan speed. Emission and excitation slit widths were set to 5 nm each, and photomultiplier voltage to 600V or 700V depending on sample emission intensity. Protein concentrations in the cuvette were typically 3 μM.

### Liposome flotation assay

Samples containing 55 μM protein in 50 mM HEPES ph 7 were incubated with liposomes for 30 min at room temperature. Sucrose was added to 40% w/v, the samples were loaded in an ultracentrifuge tube, and sucrose 25% w/v and sucrose 8% w/v (both buffered with 50mM HEPES pH 7) were layered on top, forming a discontinuous sucrose gradient. Afterwards, samples were ultracentrifuged for 1h at 4°C and 125000 g in a Beckman SW Ti90 rotor, and fractions along the gradient were analyzed by SDS-PAGE to assess the final distribution of the MBL protein.

### Determination of protein orientation in liposomes by proteolysis with proteinase K

Proteoliposome samples were incubated overnight at 45°C after addition of 30 μg/mL proteinase K and 1 mM CaCl2. The proteolysis reaction was stopped by addition of 5 mM PMSF, and samples were analyzed by SDS-PAGE.

### Coarse-grained molecular simulations

In CG-MD simulations, the enzyme was located at 4 nm at least from the bilayer, in order to avoid an early protein/membrane sensing event that might bias the subsequent interaction. The Martini2.2p (polarizable) force field ^31^ was used for all the CG-MD simulations. The crystal structure of NDM-1 and VIM-2 were taken from the Protein DataBank (PDB codes: 5ZGE ^30^ for NDM-1 and 1KO3 ^42^ for VIM-2) and turned into a coarse-grain model with the Martinize tool provided by the Martini team. All the lipid bilayers were generated with the Insane tool of Martini ^43^, with a lipid composition of 91% PEs, 6% CDLs and 3% PGs for the outer membrane, and 67% PEs, 5% CDLs, 28% PGs for the inner one, according to lipidomics analyses present in literature ^29,32^. For what concerns the acyl chains, since no reliable data are present in literature, we used the motifs that appear most frequently in nature, that are: one 16:0 and one 18:1 (1-palmitoyl-2-oleoyl) for PEs and PGs, and four 18:1 for CDLs. All the systems were solvated with the polarizable Martini water model ^44^ and ionized in 150 mM of NaCl. Each system (NDM-1 in apo/holo form; with/without post-translational modification; in presence of inner/outer bacterial membrane; with/without cardiolipins. VIM-2 in holo form; in presence of inner/outer bacterial membrane. N-VIM in apo/holo form; with/without post-translational modification; in presence of outer bacterial membrane. NDM-1 R45E-R52E in presence of outer bacterial membrane) was repeated in 5 distinct replicas. Each replica was simulated for 2 *μ*s; the systems for which we evaluated the lipid distribution were elongated to 10 *μ*s, as reported in **Table S1**. These trajectories, taken together, sum up to a total simulation time of 550 *μ*s. Frames for the analysis were collected every 750 ps. A binding event (**Table S1**) is considered to have occurred when the protein settles at ~3 Å distance from the membrane, and this distance remains constant for the rest of the simulation (i.e. no detachment occurs). For each system, the equilibration procedure was run as follows: first, the system went through 5000 steps of minimization using the steepest descent algorithm; successively, it was equilibrated with 5 ns of MD in NVT conditions, using Particle Mesh Ewald (PME) for the electrostatic contributions and velocity rescale algorithm for temperature coupling at 310 K. The production phase was conducted in NPT ensemble, using a Parrinello-Rahman semi-isotropic coupling algorithm ^45^ for maintaining the pressure constant at 1 bar. The post-translationally modified Cys26 was built using the parameters for a similar lipid present in literature ^46^. The linker between the cysteine and the lipid was modelled, following the guidelines provided by the Martini developers. The two zinc ions in the catalytic site were represented as one single Martini bead of Qa type, connected to the 6 coordinating residues (His120, His122, Asp124, His189, Cys208, His250) through harmonic potentials. Such particle was charged with +2e, that is the net charge of the catalytic site (2 zinc ions charged +2e each; 1 hydroxyl ion charged −1e; the deprotonated Cys225 charged −1e). The decision to model the two zinc ions with one single particle was taken because all the available Martini beads are characterized by a van der Waals radius that is too large to fit two particles within the limited free volume that is present in the catalytic site. The electrostatic potential generated by our model was compared to the atomistic representation and, within the limits of the CG modelling, the two are shown to have a good agreement (**Figure S3**).

### All-atom molecular simulations

Out of all the CG-MD, 3 replicas for each condition of NDM-1 (apo/holo form, with/without post-translational modification) were selected to be back-mapped into atomistic systems. In particular, the lipids were back-mapped using the Backward Martini tool ^47^. The same tool is not efficient enough to reproduce reliable atomistic representations of proteins: therefore, in order to add the enzyme to the atomistic system, we took the crystal structure available at the Protein DataBank and, for each replica, we aligned it to the CG structure of the same protein in the last frame of the CG-MD simulation. With this procedure, it was possible to preserve the lipid distribution obtained during the CG simulation, without compromising the quality of the atomistic structure of the proteins. We used Amber ff99SB-ILDN ^48^ to parameterize the protein, while the parameters for the lipids were taken from the Lipid14 ^49^ repository of Amber. Lipid14 allows to generate lipid models in a modular way, that is by combining the polar heads with appropriate acyl chains. The CDL polar head is still unavailable in Lipid14, therefore we used an Amber parameterization obtained by Lemmin et al. ^50^. The catalytic site was modelled according to a parameterization from Merz et al. ^51,52^. In this model, the charge distribution was updated for all the atoms of the six residues that coordinate the two zinc ions, to model the charge transfer due to the presence of the metal ions.

The parameters for the post-translationally modified Cys26 were generated ex-novo following the Amber guidelines. All the AA systems were solvated with a TIP3P solvent model and ionized in 150 mM NaCl. Since the atomistic systems were coming from a back-mapped CG system, the equilibration procedure had to be particularly accurate, in order to avoid clashes: initially, we followed the procedure suggested by the Martini team, which consists in performing a first minimization step after inactivating all the non-bond interactions (Coulomb and Lennard-Jones); non-bond interactions are then gradually restored in successive MD equilibration stages. After having reached a reasonable equilibrium, we performed another stage of minimization (5000 steps of steepest descent algorithm), followed by 5 different stages of MD equilibration in NVT ensemble (for a total of 500 ps), with harmonic constraints applied both on the protein backbone and on the lipids phosphates, which were gradually released. The production phase was run in NPT ensemble, using the Verlet algorithm ^53^ for the neighbor search, velocity rescale temperature coupling algorithm ^54^ for maintaining the temperature constant at about 310 K and Parrinello-Rahman pressure coupling algorithm ^45^ in semi-isotropic conditions at a pressure of 1 bar. The electrostatic interactions were treated with Particle Mesh Ewald method ^55^. Each AA MD replica was simulated for 800 ns, for a total of 9.6 *μ*s (12 replicas). Frames for analysis were collected every 100 ps.

Both in the CG and in the AA simulations, the lipid occupancy was evaluated through the VolMap Tool plugin of VMD, in particular by calculating the average occupancy of each lipid phosphate throughout the trajectory. The electrostatic potentials calculations were performed through the APBS software ^56^, which solves the Poisson-Boltzmann equations to compute the electrostatic potentials of biomolecules. The dielectric constant was set at 78 for the solvent, and 2 for the protein; the ionic concentration was set at 150 mM NaCl; protonation states of the protein aminoacids were assigned with PropKa ^57^.

## SUPPORTING INFORMATION

Output summary of CG and AA MD simulations; further analysis of lipid occupancy; electrostatic potential analysis of CG and AA models of NDM-1.

## AUTHOR INFORMATION

## Corresponding Author

* To whom corresponding should be addressed

## Author Contributions

A.P., G.B., L.J.G., A.J.V. and M.D.P. conceived the project and designed the experiments. A.P. performed the computational experiments. G.B. performed the wet lab experiments. A.P., G.B., L.J.G., A.J.V. and M.D.P. wrote the manuscript. A.J.V. and M.D.P. provided the financial funding.

## Acknowledgments

We thank Dr. Deniz Aydin for fruitful discussions. This work was supported by a MinCyT (SUIZ/17/10), Swiss National Science Foundation grant (514106) to A.J.V. and M.D.P, and grants from ANPCyT (PICT-2016-1657) and NIH (2R01AI100560-06A1) to A.J.V. G.B. is recipient of a fellowship from CONICET, and L.J.G. and A.J.V. are staff members from CONICET.

## Abbreviations

MBL: metallo-*β*-lactamase
NDM-1: New Delhi metallo-*β*-lactamase
OMV: outer membrane vesicle
CG: coarse-grain
AA: all-atom
MD: molecular dynamics
PE: phosphatidylethanolamine
PG: phosphatidylglycerol
CDL: cardiolipin

## Notes

The authors declare no competing financial interests.

## Supplementary figures

**Figure S1.**
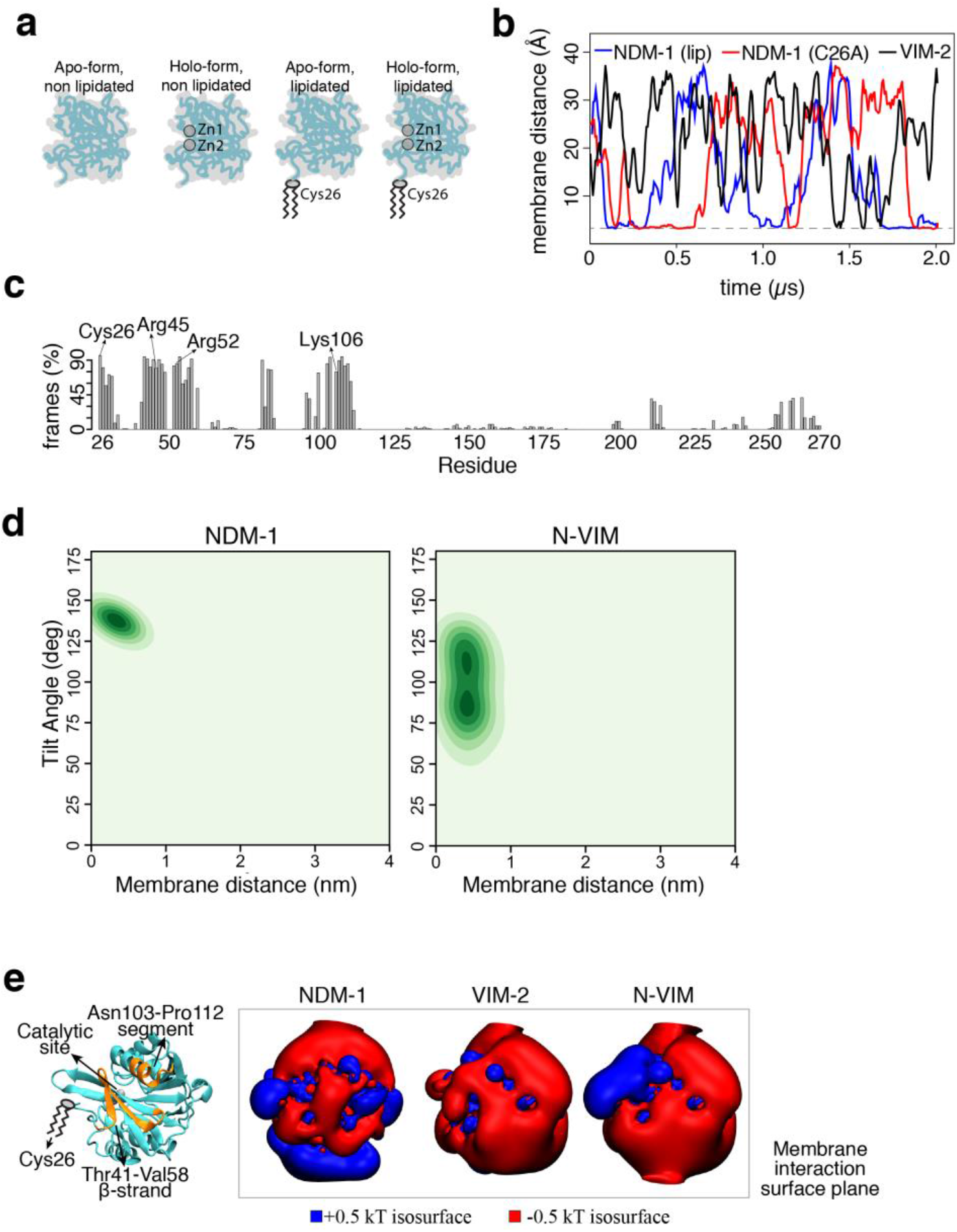
Relevant differences in membrane interaction between NDM-1, VIM-2 and N-VIM. (**a**) Summary of the protein states that were studied with CG-MD simulations (see also Table S1). (**b**) Protein-membrane distance evolution for lipidated NDM-1, soluble NDM-1 and VIM-2 in presence of the inner bacterial membrane model. One representative replica for each protein is reported. (**c**) Contacts with the membrane, expressed in percentages of total number of frames (collected every 750 ps) for each NDM-1 residue across the CG-MD simulations. (**d**) Tilt angle distribution vs. protein-membrane distance across the CG MD simulations for NDM-1 and N-VIM, respectively. (**e**) Electrostatic potential isosurfaces at ±0.5 kBT for NDM-1, VIM-2 and N-VIM, respectively.

**Figure S2.**
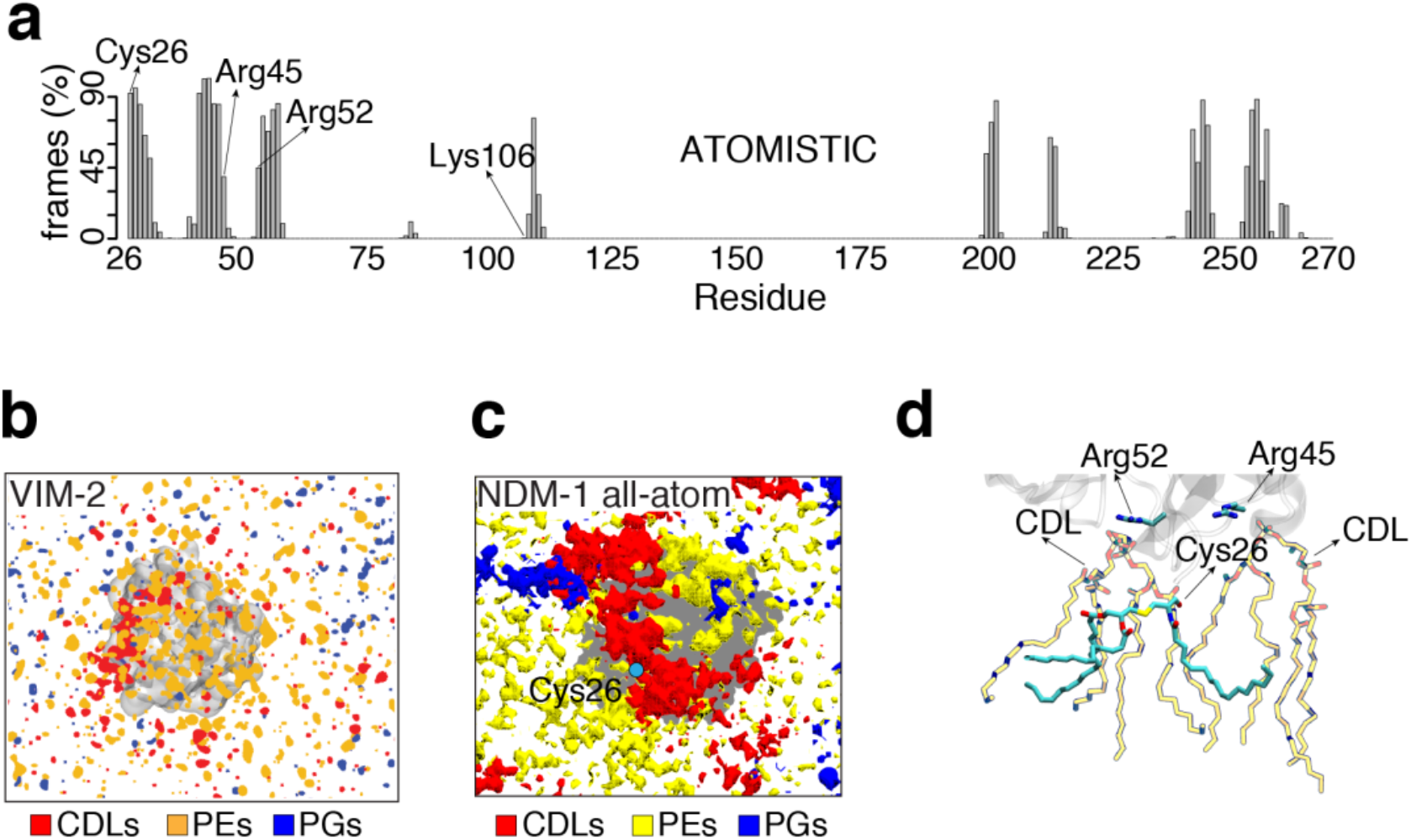
CDLs drive the interaction between NDM-1 and the outer bacterial membrane. (**a**) Contacts with the membrane, expressed in percentages of total number of frames, collected every 750 ps, for each NDM-1 residue across the AA-MD simulation. (**b**) Lipid distribution in proximity of VIM-2 during CG-MD simulations. (**c**) Lipid distribution in proximity of lipidated NDM-1 in holo-form during AA-MD simulations. (**d**) CDLs interacting to Arg45 and Arg52.

**Figure S3.**
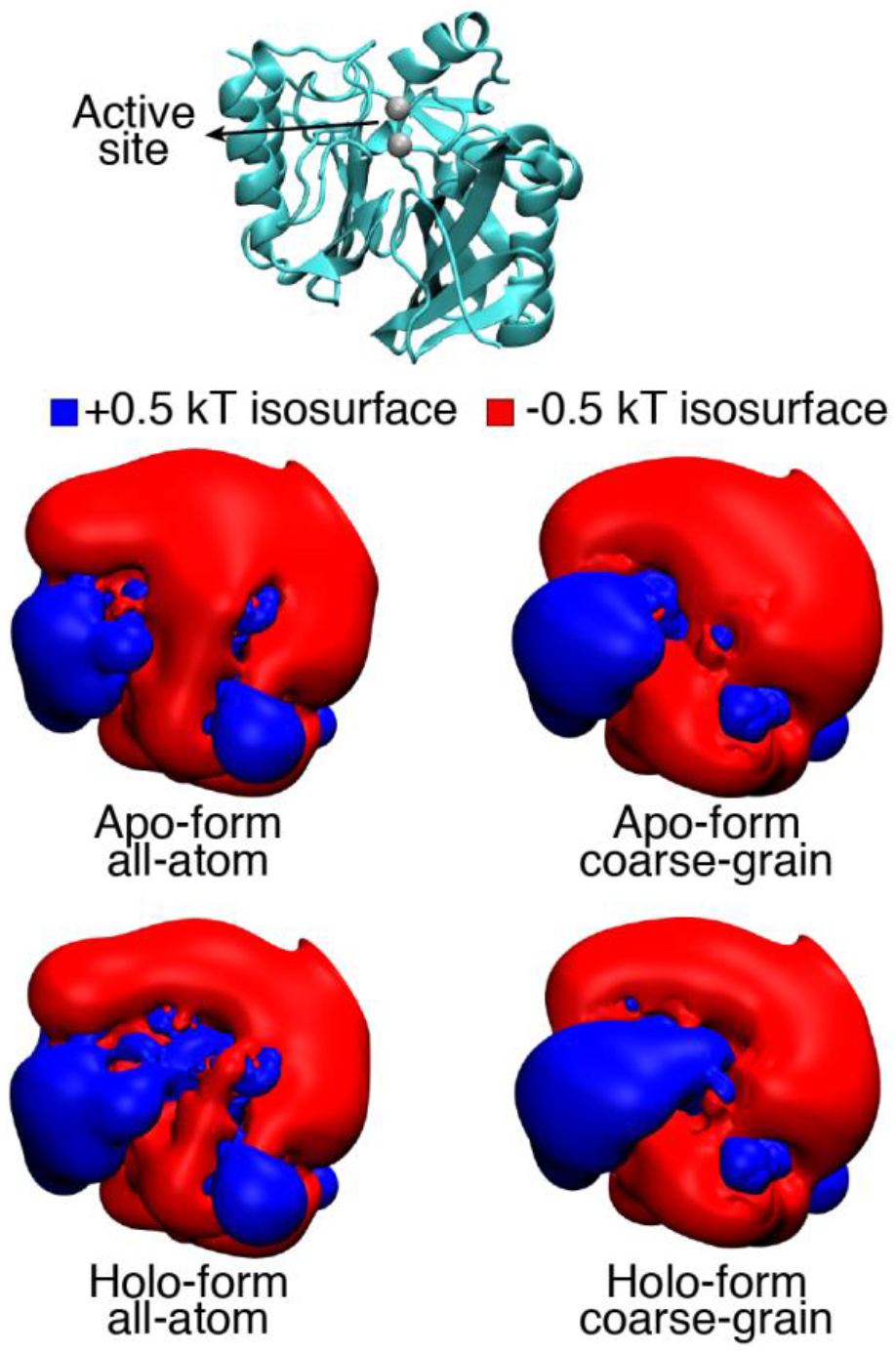
Electrostatics in NDM-1 models. Electrostatic potential isosurface for the apo/holo forms of NDM-1 in the AA and CG model.

## Supplementary table

**Table S1.**
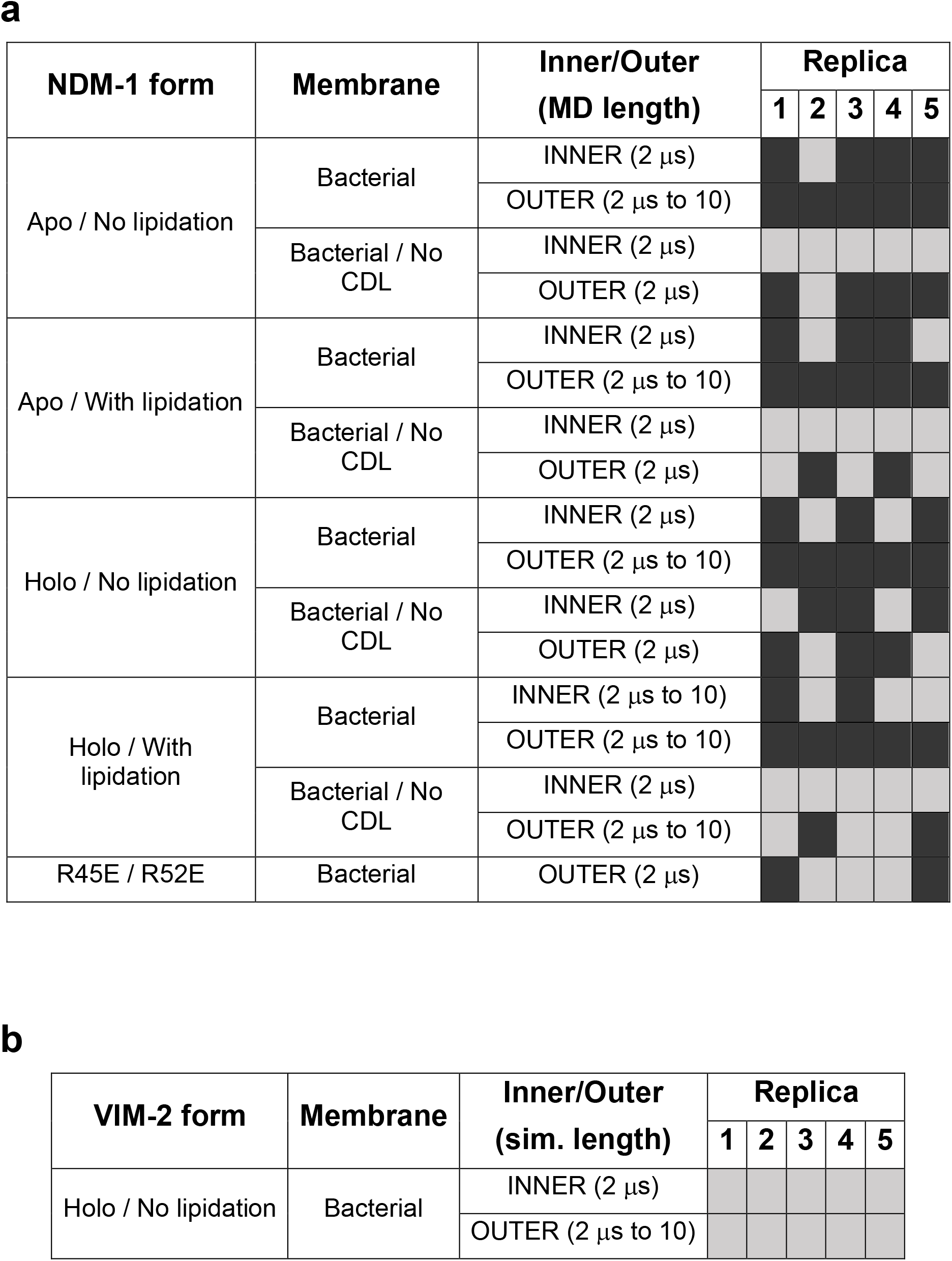
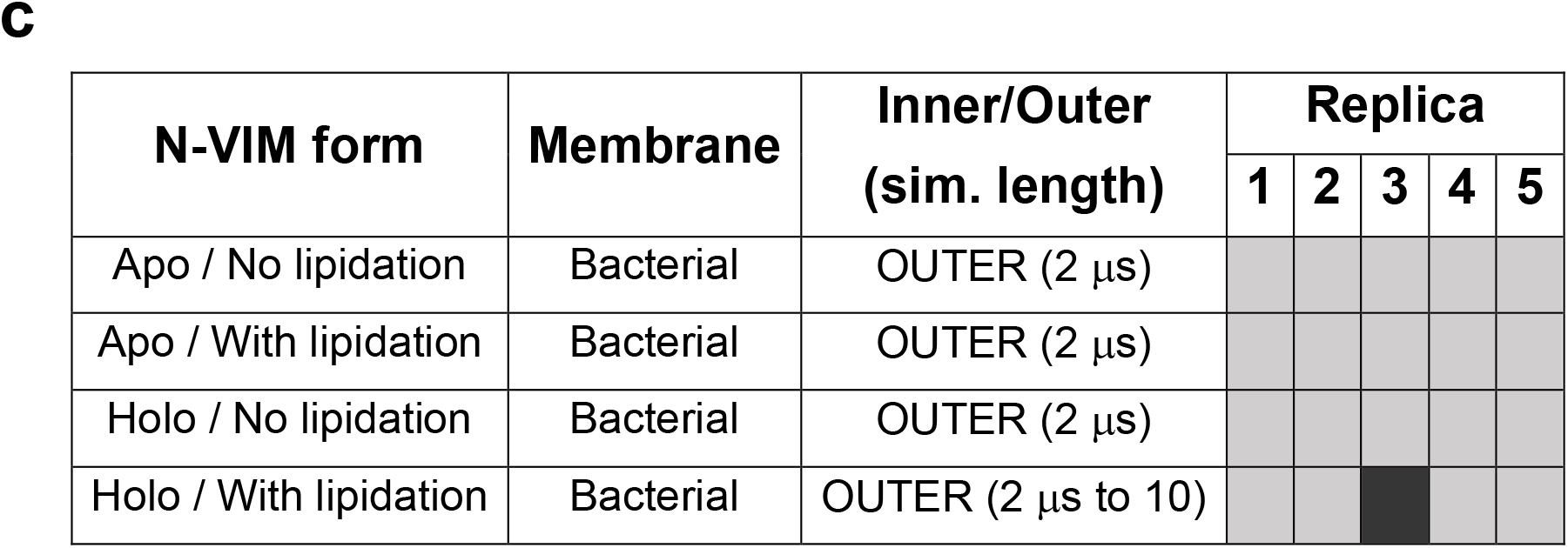
Summary of the CG-MD simulations output. List of all CG-MD simulation and the relative binding events observed for (**a**) NDM-1, (**b**) VIM-2 and (**c**) N-VIM, in different protein and membrane states. A black box means that the protein/membrane binding event has happened, while a grey box means that no binding was observed.

